# Expanding Glycopeptide Identification with Match-Between-Glycans in FragPipe

**DOI:** 10.64898/2026.02.18.706650

**Authors:** Jiechen Shen, Daniel A. Polasky, Shelley Jager, Fengchao Yu, Albert J. R. Heck, Karli R. Reiding, Alexey I. Nesvizhskii

## Abstract

Glycosylation is one of the most important, but also most complex, post-translational modifications of proteins, playing a pivotal role in various pathological processes. Mass spectrometry-based large-scale glycoproteomics analysis offers a powerful approach to explore the fundamental roles of glycosylation in both physiological and pathological contexts. Traditionally, DDA glycopeptide assignment relies on information-dense MS2 spectra, containing sufficient fragmentation information to identify both the peptide and glycan moieties. Achieving this fragmentation can be difficult, especially for low-abundant glycopeptides and/or large, complex glycans. These glycopeptides are often not assigned using current data analysis software, yet they can be of biological relevance. Here, we introduce a method called match-between-glycans (MBG), which expands glycopeptide identification while maintaining the existing glycoproteome analysis workflow. MBG enables expanding the set of identified glycopeptides to include those without MS2 spectra, or with lower quality MS2 spectra, by looking for MS1 signals displaced from other identified glycopeptides by one or multiple monosaccharide unit(s). MBG can also identify glycans not included in the glycan database, such as those containing adducts or modifications, allowing these glycans to be recovered without a drastic expansion of the search space. Combined with target-decoy FDR control, we show this method is capable of accurately expanding glycopeptide identifications and providing a more complete quantitative profile of glycosylation at each glycosite. MBG is fully integrated into the glycoproteomics workflows in FragPipe, allowing seamless, one-click operation.

## Introduction

Glycosylation is one of the most important and complex co-/post-translational modifications of proteins, regulating protein function, stability, and localization^1,2^. Mass spectrometry based glycoproteomics offers a robust way to tackle the overwhelming complexity of protein glycosylation^3-6^. Recent years have seen dramatic improvements in methods for sample preparation and mass spectrometry of glycopeptides^7-9^. More notably, the recent proliferation of glycoproteomics analysis tools significantly enhances the depth and breadth of glycoproteome studies across diverse applications^10-20^. However, due to the stochasticity in MS/MS acquisition, low abundance glycopeptides are often missed, either because no MS2 spectrum was acquired or due to low spectral quality. This issue is much more prevalent in glycoproteomics than in standard proteomics as the various co-occurring glycans that are attached to the same peptide, or glycosylation site, dilute the ion signal for each individual glycopeptide. This poses a significant challenge for comprehensive glycopeptide identification using traditional data-dependent acquisition (DDA) methods^6^. This is particularly problematic when investigating changes in glycosylation across large sample cohorts, as real biological variation becomes difficult to distinguish from differences arising from stochastic sampling.

Protein glycosylation is a non-template-driven, stepwise biosynthetic process in which mono- and polysaccharides are sequentially added to (and removed from) growing glycan chains. As a result, the glycans attached to the same peptide backbone often form a series of closely related structures that differ by one or a few monosaccharide units^2^. The resulting range of glycopeptides sharing the same peptide sequence but differing in their attached glycan structures are known as glycoforms. In reverse-phase liquid chromatography, the retention behavior of glycopeptides is primarily governed by the peptide backbone rather than the glycan moiety, aside from a few notable exceptions, due to the stronger hydrophobic interactions of amino acid side chains with the stationary phase^21^. As a result, glycopeptides sharing the same peptide backbone tend to elute within a narrow retention time (RT) window, except for glycopeptides with sialic acids, which significantly delay RT^21,22^. Given this, and using the mass difference between monosaccharides, several methods have previously been developed to attempt to expand the number of glycoforms identified for a glycopeptide. Utilizing the features of retention time change, Klein et al. developed a linear modeling approach to correcting the identifications from public datasets^23^, and applied similar concept for PSM rescoring to boost identification^24^. Nagai-Okatani et al. proposed a MS1-based method Glyco-RIDGE for inferring glycoforms, which was further developed as the software GRable for automating this analysis^25,26^. However, the GRable workflow relies on RT revision from deglycosylated peptides, complicating sample preparation, and only provides a built-in “confidence level” instead of an FDR or q-value, posing an issue for larger-scale applications.

Here, we introduce an MS1-based method called match-between-glycans (MBG), a lightweight, modular method designed to expand glycopeptide identification by leveraging learned RT and/or IM shifts between related glycan compositions. MBG starts from high-confidence glycopeptide identifications and systematically infers neighboring glycoforms based on well-known and well-defined glycan or adduct mass differences. These potential glycoforms are searched against MS1 features, and inferred targets are statistically evaluated using a decoy-based scoring model before being integrated back into downstream pipelines.

We validated MBG across a series of diverse datasets. In a benchmark analysis using a fission yeast dataset, where most N-glycans are of the high-mannose type, MBG recovered many expected glycoforms with high RT consistency, leading to a 23.6% increase in glycopeptide identifications (peptide + glycan combinations) at 5% FDR. Applied to human plasma glycoproteomics data set acquired by PASEF, MBG identified additional sialylated glycoforms, improving site-specific glycan coverage. We also applied MBG in a TMT-labeled glioblastoma (GBM) dataset from the CPTAC project, combined with FragPipe-Analyst^27^, to demonstrate the utility of MBG for isobaric labeling quantification of glycoproteomics data. Finally, MBG was shown to efficiently identify glycopeptides with adducts (e.g., NH_4_^+^, Fe^3+^, Na^+^) and even more rare glycan modifications such as mannose-6-phosphate (M6P), without requiring prior database inclusion or extensive search space expansion. Together, these results establish MBG as a robust and accessible strategy for expanding glycopeptide coverage. Seamlessly integrated within the FragPipe ecosystem, MBG empowers sensitive, biologically informative glycoproteomic analysis with minimal additional computational burden.

## Experimental Procedures

### Materials and sample preparation

Human plasma sample set: Human pooled plasma samples (VisuCon-F Normal Donor Set (EFNCP0125), Affinity Biologicals) were used and prepared as described previously^28^. Cotton-HILIC based enrichment of glycopeptides was performed on 6 technical replicates of trypsin plus LysC digested plasma samples. Briefly, samples were denatured, reduced and alkylated using SDC, TCEP and CAA, followed by overnight digestion with trypsin and LysC. Peptides were desalted using Oasis PRiME HLB (10 mg) 96-well plates (Waters) and enriched using 10 mm cotton thread on in-house made SPE tips. Samples were dried and resuspended in 0.1% TFA prior to LC-MS/MS analysis. Data was acquired on a timsTOF HT Mass Spectrometer (Bruker) coupled online to an Ultimate3000 UHPLC system (Thermo Fisher Scientific).

Glycopeptides were separated using an Aurora Ultimate XT (25 cm x 75 µm, C18 stationary phase) analytical column, heated to 50 °C with a constant flowrate of 400 nl/min. Peptides were loaded using 1% buffer B for 13 min, followed by a linear gradient from 3% to 35.2% B in 130 min, a column wash at 80% B for 8 min, and equilibration at 1% B for 12 min. Buffer A was 0.1% FA in water, buffer B was 0.1% FA in ACN. The timsTOF HT was operated in positive mode, with a scan range of 100-1700 *m/z*, and a tims range of 0.5-1.6 Vs/cm^2^. For MS/MS, the settings were the following: PASEF ramps, 7; charge range, 2-5, PASEF precursor region, polygon with following coordinates (264.6, 0.58), (432.31, 0.90), (1244.80, 1.47), (1700, 1.6), (1700, 0.58); stepping collision energy (SCE), 35 and 40 eV at 0.50 Vs/cm^2^ and 65 and 100 eV at 1.6 Vs/cm^2^, as was described before^29^.

Other Dataset Details: “Yeast”, “mouse liver” and “mouse brain” datasets were downloaded from Proteome Xchange^30^ repositories with identifier PXD005565, PXD005553 and PXD005411, respectively. The raw data was converted to mzML with MSConvert^31^. Glycopeptides of these three datasets were enriched by ZIC-HILIC and analyzed with Orbitrap Fusion mass spectrometer using stepped-energy (NCE=20-30-40) HCD for fragmentation. The GBM dataset was downloaded from Proteomic Data Commons (PDC) at: https://pdc.cancer.gov/. Mixed anion exchange (MAX) enriched glycopeptides were labeled with 11-plex TMT reagents and analyzed by Orbitrap Fusion Lumos Tribrid with HCD with a normalized collision energy (NCE) of 35^32^.

### Data Analysis Using MSFragger-Glyco and Match-Between-Glycans (MBG)

The “Yeast” glycoproteomics data was searched against a yeast proteome database includes common contaminant proteins and decoys (downloaded 07/02/2025, 5,236 non-decoy entries). A yeast and mouse combined glycan database with 1701 entries (containing the glycans from Mouse_N-glycans-1670-pGlyco in addition to high-mannose glycans ranging from HexNAc(2)Hex(3) to HexNAc(2)Hex(20)) was used for yeast glycan assignments, with a maximum of one ammonium adduct allowed per glycan, generating a total of 2315 unique glycan masses. The digestion enzyme was set as trypsin with up to two missed cleavages allowed. Variable modifications included methionine oxidation and protein N-terminal acetylation. Searches were run in N-glycan mode with precursor and fragment mass tolerances of 20 ppm and 10 ppm, respectively. Isotope error correction was enabled, and ion types b, y, b + HexNAc, y + HexNAc, and Y were considered. Oxonium ion filtering was also applied using common oxonium masses, requiring a minimum relative intensity of 10% to consider spectra for glycopeptide identification. Philosopher (v5.1.1)^33^ was used for peptide-level FDR filtering. PSM probabilities were modeled using PeptideProphet^34^ in semi-parametric mode with the extended mass model (mass width: 4000), N-glycan motif modeling enabled, and cLevel set to 0. Protein inference was performed with ProteinProphet^35^ using default parameters, except maxppmdiff was set to 20,000,000 to retain glycopeptides with large delta masses. Search results were then subjected to a set of MBG searches including an entrapment search. FDR thresholds were set to 1% at the PSM, ion, peptide, and protein levels. PTM-Shepherd was enabled for glycan assignment and glycan FDR filtering before conducting MBG searches^14,36^. The parameters for MBG searches were as follows: maximum glycan shift steps – 2, maximum q-values – 0.01, minimum glycans per sites – 2, minimum PSMs – 1, RT tolerance – 0.4 minutes, m/z tolerance – 10 ppm. FDR was set as 0.01 and 0.05 respectively. Glycan composition shift was set as “Hex(1), NH4(1)” for normal search and “Hex(1), NH4(1), NeuAc(1), HexNAc(1)Hex(1)NeuAc(1)” for entrapment search.

The “Human plasma PASEF” glycoproteomics data was searched against a human proteome database that included common contaminant proteins and decoys (downloaded 07/02/2025, 20,453 non-decoy entries). “Human N-glycans medium-253” was used for the Glycan database, which contained 506 glycans after allowing a maximum of one NH4 per glycan. “IM-MS” was checked as MS data type. For the MBG search, allowed glycan composition shift was set as “HexNAc, Hex, Fuc, NH4, NeuAc, HexNAc(1)Hex(1)NeuAc(1), HexNAc(1)Hex(1)”. Other parameters were the same as used for the yeast data searching.

“GBM” data was searched against the same protein database as Clinical Proteomic Tumor Analysis Consortium (CPTAC) glioblastoma (GBM) paper^32^, but a glycan database that did not include any adducts. The “glyco-N-TMT” workflow was used for searching with label type set as TMT-11. MBG was performed with the same parameters except for glycan composition shift was set as “HexNAc, Hex, Fuc, NeuAc, HexNAc(1)Hex(1)NeuAc(1), HexNAc(1)Hex(1)”. Other parameters were the same as yeast data searching.

“Mouse liver” and “mouse brain” data were searched against a mouse proteome database including common contaminant proteins and decoys (downloaded 07/03/2025, 17,352 non-decoy entries). The same glycan database as “yeast” data search was used for “mouse liver” data search. For “mouse brain” data search, to consider mannose-6-phosphate (M6P) glycans, a small database with 196 entries was used for M6P glycan search, which included 14 phospho-glycans. For MBG search, glycan composition shift was “HexNAc, Hex, Fuc, NeuAc, HexNAc(1)Hex(1)NeuAc(1), HexNAc(1)Hex(1), Fe, Na, NH4” for “mouse liver” data search and “HexNAc, Hex, Fuc, HexNAc(1)Hex(1), Phospho, NH4” for “mouse brain” data search.

Downstream analysis was performed with FragPipe-Analyst and visualization with FragPipe-PDV viewer.

### Parameters of MBG

MBG FDR: FDR filtering for MBG inferred glycopeptides, which is based on the precursor level. Maximum glycan shift steps:

Once an inferred glycopeptide is matched by IonQuant, the MBG process of iteratively testing additional mono or polysaccharide units continues based on that matched glycopeptide. This parameter defines the maximum number of MBG process steps permitted per inference chain. Note that chains only continue up to this maximum if a glycoform is matched at each intermediate step. A value of 0 disables MBG entirely, preventing any inference.

Maximum Glycan Q-value, Minimum Glycans, and Minimum PSMs:

These parameters define the filtering criteria for glycopeptides eligible for MBG processing. A glycopeptide must meet all three conditions to proceed: (1) it must be supported by at least the specified number of PSMs (“Minimum PSMs”) (2) each supporting PSM must have a glycan q-value below the “Maximum Glycan Q-value” threshold, and (3) the corresponding glycosite must be associated with at least the specified number of distinct glycans (“Minimum Glycans”).

Glycan composition shift: monosaccharide or a set of monosaccharides (or any glycan modifications/adducts) shifts used for MBG inferring from confidently identified matches. Parameters of matching peaks in MS1 can be modified in “Quant (MS1)” panel of FragPipe, including *m/z*, RT and IM tolerance.

## Results and Discussion

Match-between-glycans, or MBG, is an independent method of expanding the identification of glycopeptides following a database search method, which leaves the initial search unchanged. As shown in **Fig. 1**, the MBG workflow begins with confidently assigned glycopeptides from MSFragger-Glyco search combined with glycan assignment of PTM-Shepherd. Parent PSMs are first selected based on user-defined criteria, including minimum PSM count, minimum number of glycans per peptide, and glycan q-value thresholds. For each selected parent PSM, MBG then generates alternative glycoforms that differ from the originally identified glycan by one monosaccharide or a user-defined set of monosaccharides. The masses of these potential glycoforms are then searched in nearby MS1 spectra using IonQuant^37,38^. The expected retention time (RT) and/or ion mobility (IM) of the MBG glycopeptides are based on median shifts observed for analogous glycan changes observed in the dataset, and candidate matches are evaluated within a user-defined tolerance window. The variability of the shift distribution is not explicitly incorporated into the prediction in favor of the user-defined tolerance to prevent large tolerances in the case of outliers for shifts with relatively few contributing PSMs. More specifically, two tables are generated during the process: one is a glycan shifts table, which is comprised of RT/IM shifts between specific glycans, like HexNAc(2)Hex(8) to HexNAc(2)Hex(9), and is given priority in usage. Alternatively, a unit shift table records median shifts of monosaccharide changes (e.g., Hex(n) to Hex(n+1) on any glycan), which is used when there is not a corresponding shift in the specific glycan shift table. Meanwhile, for each target glycoform searched, MBG generates one decoy glycoform that shares the same RT and IM but has an 11 Da increment in mass. 11 Da was selected to avoid common isotopes and modifications of the initial glycopeptide, which could confound the decoy process. All identified targets and decoys are subsequently subjected to LDA modeling for scoring. After the FDR filtering, the inferred glycopeptides are written back to the psm.tsv file for downstream processing in FragPipe. Seven features are considered for scoring: RT and/or IM shift, mass error, precursor intensity, relative intensity of the Y0 and Y1 ions (for those glycoforms that have corresponding MS2 spectra found), Kullback-Leibler divergence (an isotopic envelope shape match score from IonQuant) and frequency of the glycan shift (from identified PSMs). See the Methods section for the detailed MBG parameters used in each search. It is worth noting that MBG controls FDR at the precursor level, as it is fundamentally an MS1-based method, ensuring rigor in glycopeptide inference without inflating identification counts.

**Figure 1.**
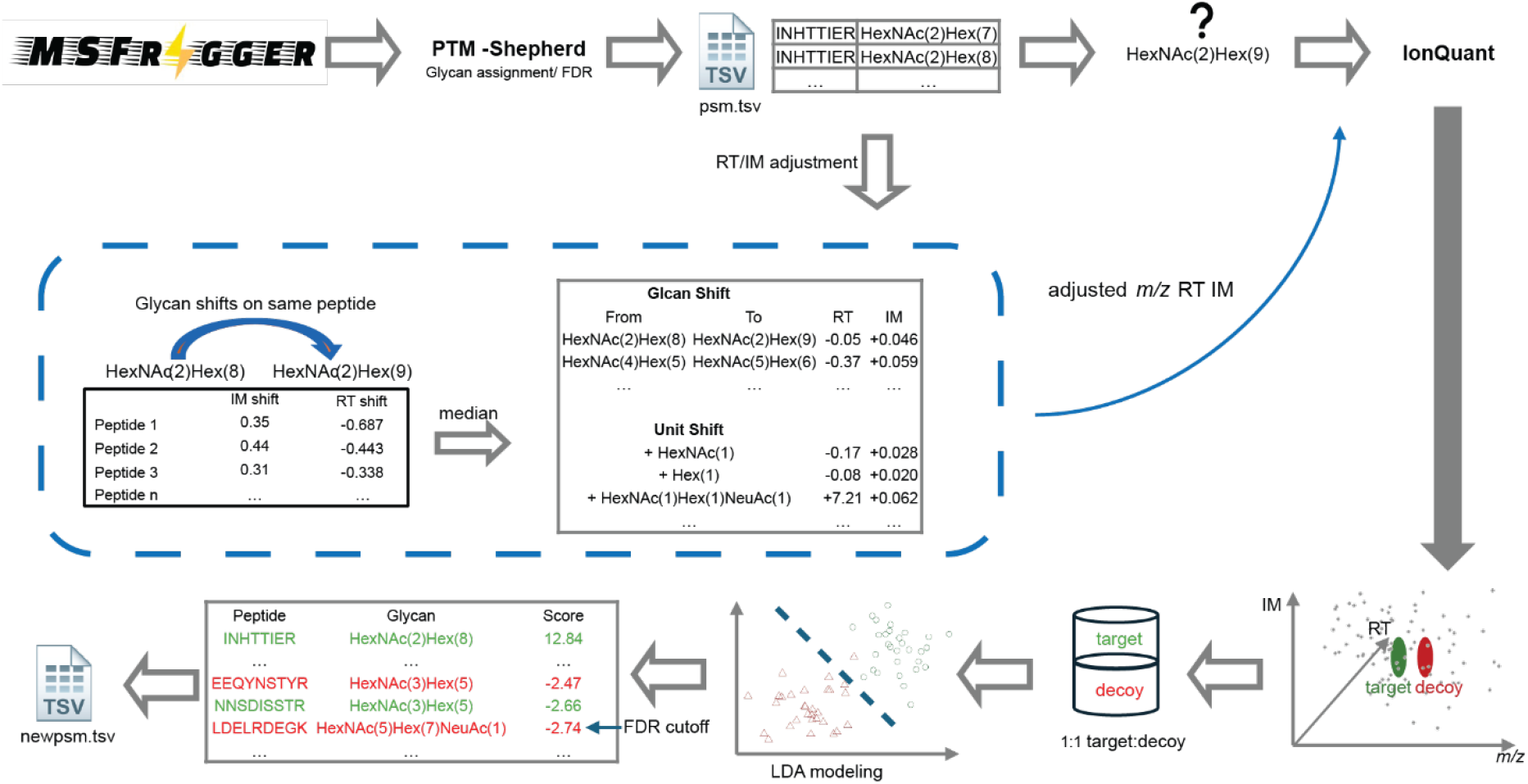
Workflow of MBG. MBG operates after MSFragger-Glyco and PTM-Shepherd. For each confidently assigned glycopeptide that passes user-defined filters, MBG generates alternative glycoforms differing by one monosaccharide (or a defined group of monosaccharides) and searches for their precursor masses within expected retention time (RT) and/or ion mobility (IM) windows using IonQuant, with these windows predicted from empirically observed glycan shifts or, when unavailable, monosaccharide unit shifts. Each target glycoform is paired with a decoy of identical RT/IM and a +11 Da mass offset. Target and decoy matches are scored using an LDA model based on seven features (detailed in Method section), and peak-level FDR filtering yields the final inferred glycopeptides, which are written back to the psm.tsv file for downstream FragPipe analysis.

### Benchmarking with Fission Yeast Dataset

As a proof of concept, we first benchmarked MBG using a fission yeast dataset—a well-characterized sample commonly employed for evaluating glycoproteomics tools^10,11,13,14,16,17^. Fission yeast produces a relatively simple and homogeneous N-glycoproteome dominated by high-mannose–type glycans, and contains far fewer proteins than mammalian systems, making it well suited for controlled methodological benchmarking. In particular, this dataset allowed us to test the core assumption underlying MBG: that RT shifts associated with glycan composition changes are relatively consistent and largely independent of the peptide backbone.

To validate this, we analyzed RT shifts associated with glycan changes in identifications from MSFragger-Glyco, where most N-glycans in fission yeast are of the high-mannose type. **Fig. 2a** shows RT shift distributions for the five most common glycan transitions. Most transitions between high-mannose glycans fall within 1 minute, and 73.2% to 78.4% of transitions are within a 0.4-minute deviation when using the median RT shift for RT prediction. When aggregating all Hex-based glycan shifts, 72.8% of PSMs fell within this 0.4-minute RT window (**Fig. 2a bottom**), showing that indeed the RT shifts are consistent and sufficiently independent of the peptide backbone to be used as a guide for MBG searching.

**Figure 2.**
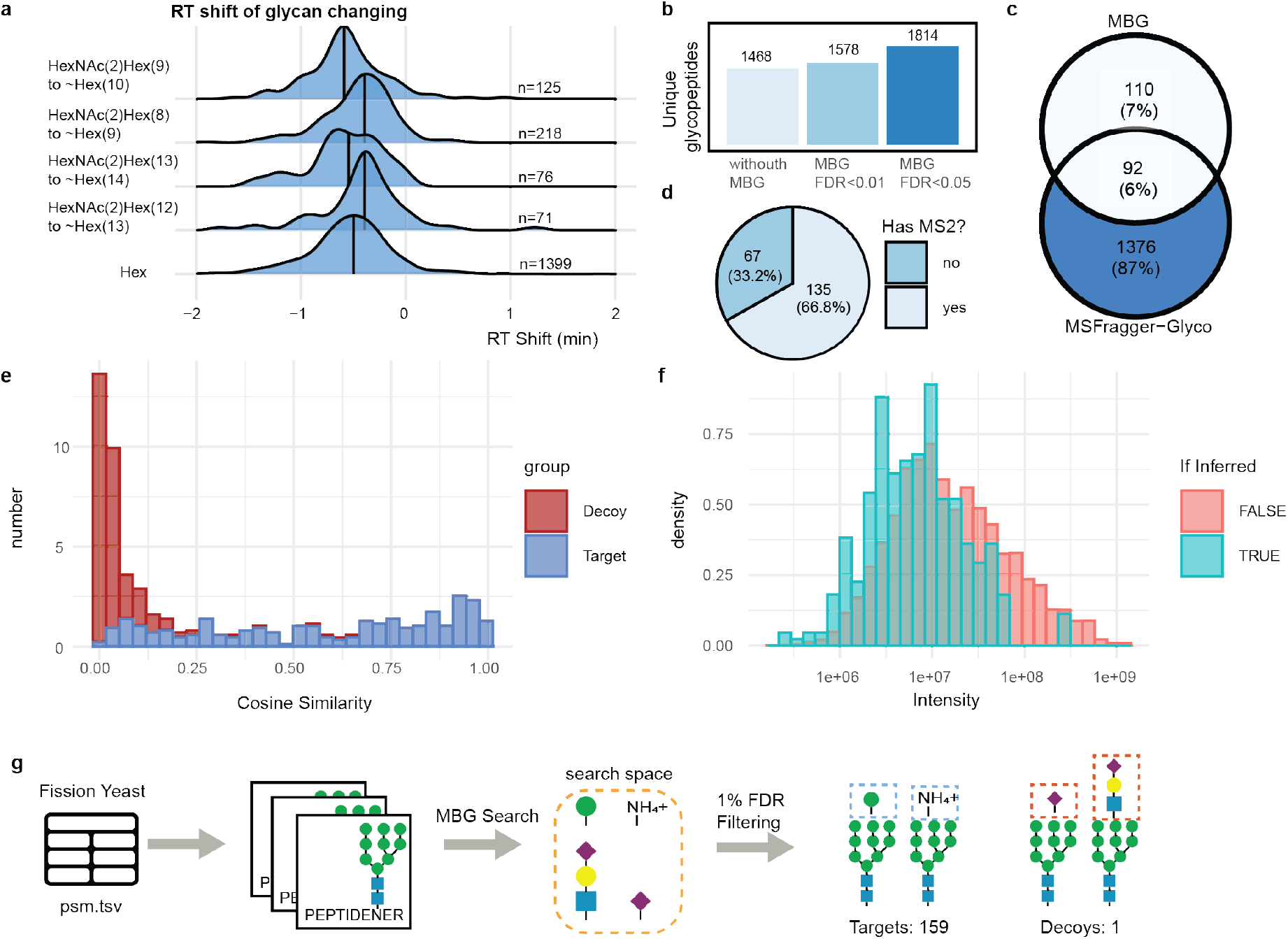
Benchmarking MBG using the fission yeast dataset. (a) RT shift distributions for the top five glycan transitions across peptides identified by MSFragger-Glyco. The black vertical line in each ridge represents the median shift of each category. (b) Identified unique glycopeptides of MSFragger-Glyco and MSFragger-Glyco + MBG at FDR thresholds of <0.01 and <0.05, respectively. (c) Overlap between MBG-inferred glycopeptides and those identified by MSFragger-Glyco. (d) Proportion of MBG-inferred glycopeptides with and without corresponding MS2 spectra. (e) Spectral similarity of inferred glycopeptides to their parent spectra compared with random spectra controls. (f) Intensity distributions of MBG-inferred glycopeptides versus those identified by MSFragger-Glyco (g) Entrapment search of MBG by including both target (Hex, NH_4_^+^) and decoy (NeuAc, HexNAcHexNeuAc) shifts into search space.

Next, we assessed the gain obtained by MBG. Applying MBG led to the identification of 7.5% and 23.6% more unique glycopeptides at FDR thresholds of 1% and 5%, respectively (**Fig. 2b**). Among the 202 glycopeptides identified by MBG at <1% FDR, 92 were also identified by MSFragger-Glyco from at least another run (**Fig. 2c**), and 135 were supported by at least one MS2 spectrum (**Fig. 2d**). Together, these results illustrate that MBG not only identifies novel glycopeptides, but also recovers glycopeptides that are observed in other runs yet missing in individual MSFragger-Glyco searches, thereby reducing missing values across runs.

To assess inference accuracy, we evaluated spectral similarity between inferred glycopeptides and their parent spectra. For comparison, we also computed cosine similarities between the same parent spectra and randomly selected glycopeptide spectra with matching charge states, similar masses, and retention times (**Fig. 2e**). MBG-inferred glycopeptides showed significantly higher spectral similarity to their parent spectra than the random control. MBG also recovered glycopeptides with lower intensity distributions compared to those identified by MSFragger-Glyco (**Fig. 2f**), demonstrating its ability to recover low-abundance glycopeptides missed by conventional searches, for example, due to the precursor ion not being selected for fragmentation.

To further assess MBG’s precision, we performed an entrapment search. Two biologically plausible shifts (Hex(1) and an NH_4_^+^ adduct) and two decoy shifts (NeuAc(1) and HexNAc(1)Hex(1)NeuAc(1), which are rare or absent in fission yeast) were included in the MBG search space (**Fig. 2g**). After applying a 1% FDR filter, only one glycopeptide containing NeuAc was identified out of 160 inferred glycopeptides (0.63%), underscoring the accuracy of MBG.

Finally, to estimate MBG’s false negative rate—i.e., glycopeptides that should be inferred but are missed—we removed all glycopeptides containing HexNAc(2)Hex(9) when the same peptide was also identified with HexNAc(2)Hex(10), and reanalyzed the data. MBG successfully recovered 68 out of 77 removed glycopeptides, corresponding to a false negative rate of 11.6%. Given that less evidence is typically available for MBG-based identifications than MS2-based database search IDs, the relatively conservative nature of MBG, with a lower false positive rate than false negative rate, is intentional to avoid spurious identifications.

### Expanding glycopeptide IDs in a Human Plasma PASEF dataset

In contrast to yeast, where N-glycosylation is limited primarily to high-mannose type glycans differing by hexoses, N-glycosylation in human plasma is far more complex. Plasma proteins are predominantly decorated with complex-type N-glycans, typically containing two to four antennae that can be capped with up to four sialic acids. These branches may be further elongated by repeating HexNAc–Hex units and fucose residues can be attached to the HexNAc moieties, adding further structural diversity. This extensive heterogeneity, combined with the high dynamic range of plasma proteins, dilutes glycopeptide signals and complicates glycopeptide identification, often resulting in incomplete coverage of the microheterogeneity at many glycosylation sites. Despite these challenges, plasma remains a highly valuable resource for biomarker discovery, underscoring the importance of comprehensive characterization of its glycosylation patterns. To this end, we applied MBG to a glycopeptide enriched human plasma DDA-PASEF dataset with the goal of improving our coverage of the glycosylation that is in the sample. The retention time shifts for most of the monosaccharide differences were normally distributed with small peak widths, with the exception of NeuAc (**Fig 3a**). NeuAc is known to cause a large retention time shift due to ionic interactions^39^. This retention time shift has a wide distribution across the different features, indicating it may be influenced by the peptide sequence and/or by the differences in sialic acid linking position, both of which are known to influence the elution profile of sialylated species^40,41^. On the other hand, the ion mobility shifts between different glycan species are normally distributed for all monosaccharide shifts: 76.2–89.9% of monosaccharide/adduct shifts fell within 0.05 IM units of the median IM shift for the same change (**Fig. 3b**). Notably, even though the NeuAc shift spans a wide retention time range, the corresponding IM shifts remain tightly distributed.

**Figure 3.**
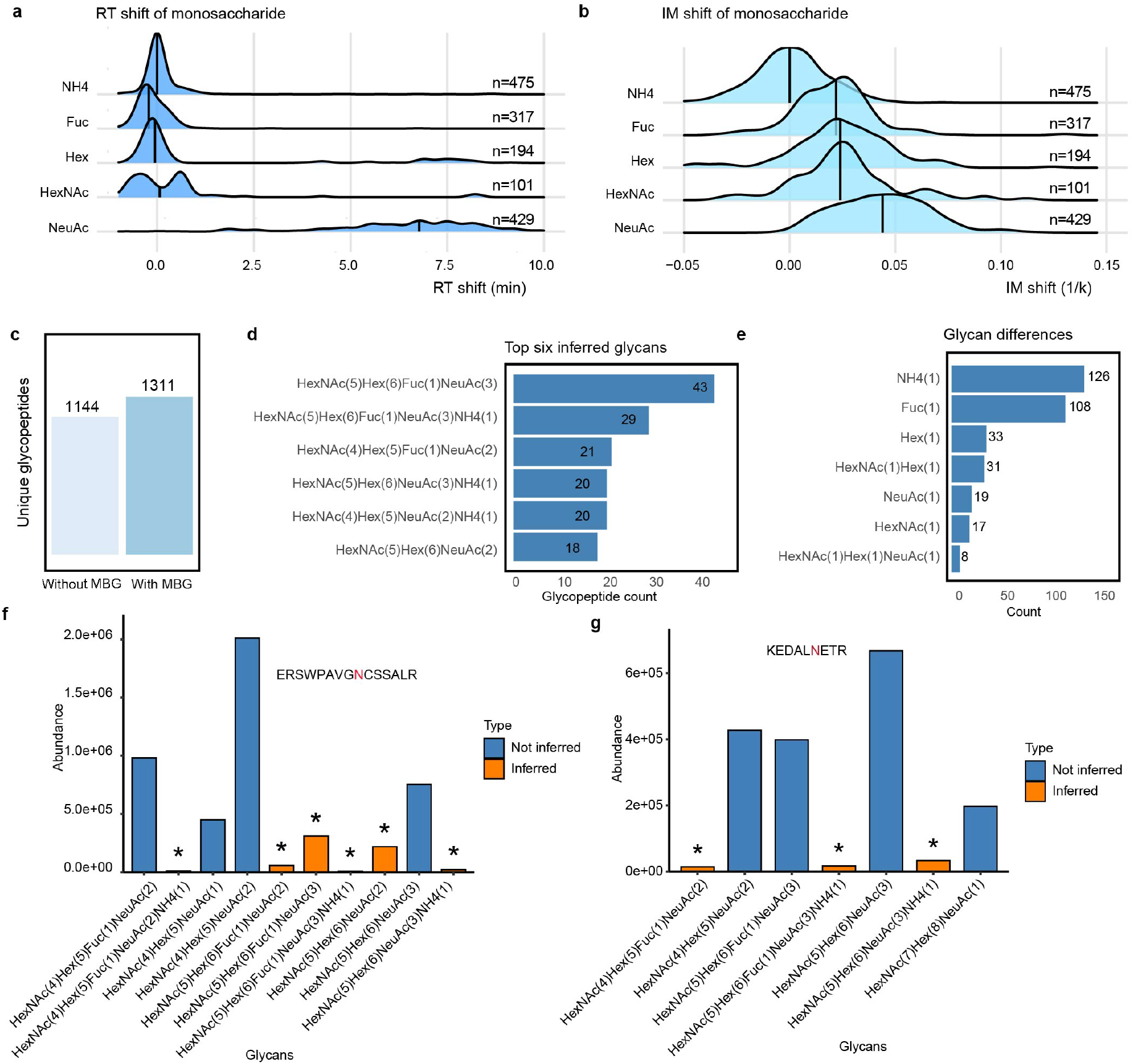
Application of MBG to the human serum PASEF dataset. RT (a) and IM (b) shifts comparison of four monosaccharide changes and an NH_4_^+^ adduct. The black vertical line in each ridge represents median shift of each category. (c) Unique glycopeptides identified with and without MBG. (d) Top six inferred glycans with MBG, counting by glycopeptide. (e) Frequency of glycan shifts between MBG inferred glycopeptides and their parent glycopeptides. Glycan landscapes and intensities of glycopeptides ERSWPAVGNCSSALR (f) and KEDALNETR (g), from which MBG restored 9 low-abundance site-specific glycans, marked with asterisks.

Across six technical replicates, MSFragger-Glyco identified 1,144 unique glycopeptides. With MBG, this number increased 14.6% to 1,311 (**Fig. 3c**). The top six inferred glycans were all sialylated complex glycans, consistent with known features of the plasma glycoproteome (**Fig. 3d**). The most frequently observed shift was the addition of the NH_4_^+^ ion, closely followed by the addition of fucose moieties (Fig. 3e). Notably, increased fucosylation of plasma proteins (such as alpha-1-antitrypsin, haptoglobin, and ceruloplasmin) has been proposed as potential biomarker across various disease states, underscoring the critical importance of accurately quantifying even low-abundant fucosylated glycan species. This is particularly pertinent because many disease-associated glycosylation changes appear in proteins present at low concentrations in plasma, and if these species are not detected, valuable biomarker signals may be missed^42-44^.

Another frequent glycan shift was HexNAc(1)Hex(1), a LacNAc moiety, whereas shifts corresponding solely to HexNAc(1) were rare. This disparity is logical: in human plasma, free HexNAc rarely exists and as it is usually galactosylated^45^, except for bisected GlcNAc. Yet bisected GlcNAc is uncommon on most plasma proteins, except those derived from B cells (e.g., immunoglobulins^46^). Notably, a sialic acid shift occurred the least frequently. This may be the result of the large spread observed in retention time shift, but might also be due to having fewer parent glycans containing sialic acid(s). We then categorized these differences by the glycan type of their parent glycans (i.e., the glycans from which MBG inferred). This showed that complex-type glycans made up the vast majority of parent glycans for almost all difference categories, with the exception of a small number of high mannose and hybrid types, which were most notable in the Hex(1) difference group (**Figure S1**).

A key strength of MBG is its ability to recover missing glycoforms at individual glycosites, providing a more complete view of the site-specific glycosylation landscape, especially on low-abundant proteins. In the plasma dataset, MBG recovered up to six missing glycans at a single glycosite. **Fig. 3f,3g** illustrates glycan landscapes for two representative sites, where MBG restored nine lower abundant glycoforms. We also compared the number of MBG-inferred glycans to their corresponding peptide intensity (**Figure S2**). This analysis revealed that peptides with higher intensity tend to be, as expected, associated with more inferred glycans. This suggests that MBG is primarily recovering low-abundance glycans from high-abundance peptides—which naturally provide more precursors for inference—rather than recovering glycans from peptides that are already low-abundance.

### Expanding glycopeptide identification from the TMT-based GBM dataset from the CPTAC project

Leveraging the extensive capabilities of FragPipe, MBG is compatible with both label-free and isobaric-labeled data. We applied MBG to reanalyze the CPTAC glioblastoma multiforme (GBM) dataset, which profiled human brain tissue samples including grade 4 astrocytomas and non-tumor controls. Previous analysis of this dataset reported widespread alterations in N-glycosylation, characterized by downregulation of fucosylated glycans and upregulation of sialylated glycans. Notably, for isobaric labeling searches, MBG ignores inferred glycopeptides that lack corresponding MS2 spectra, as these could not contribute to the quantitative output without an MS2 spectrum from which to obtain reporter ion quantities. MBG identified 5,008 glycopeptide peaks across all MS files, of which 1,005 (20.1%) were ignored due to the absence of corresponding MS2 spectra (**Fig. 4a**). After filtering, we identified 16,236 unique glycopeptides, including 14,178 (87.3%) glycopeptides exclusively identified with MSFragger-Glyco, 740 (4.6%) glycopeptides exclusively inferred by MBG and 1,318 (8.1%) inferred by MBG but also identified with MSFragger-Glyco in other runs (**Fig. 4b**). Thus, despite requiring MS2 spectra for inferred glycopeptides, many were still recovered by MBG, improving the completeness of the result set.

**Figure 4.**
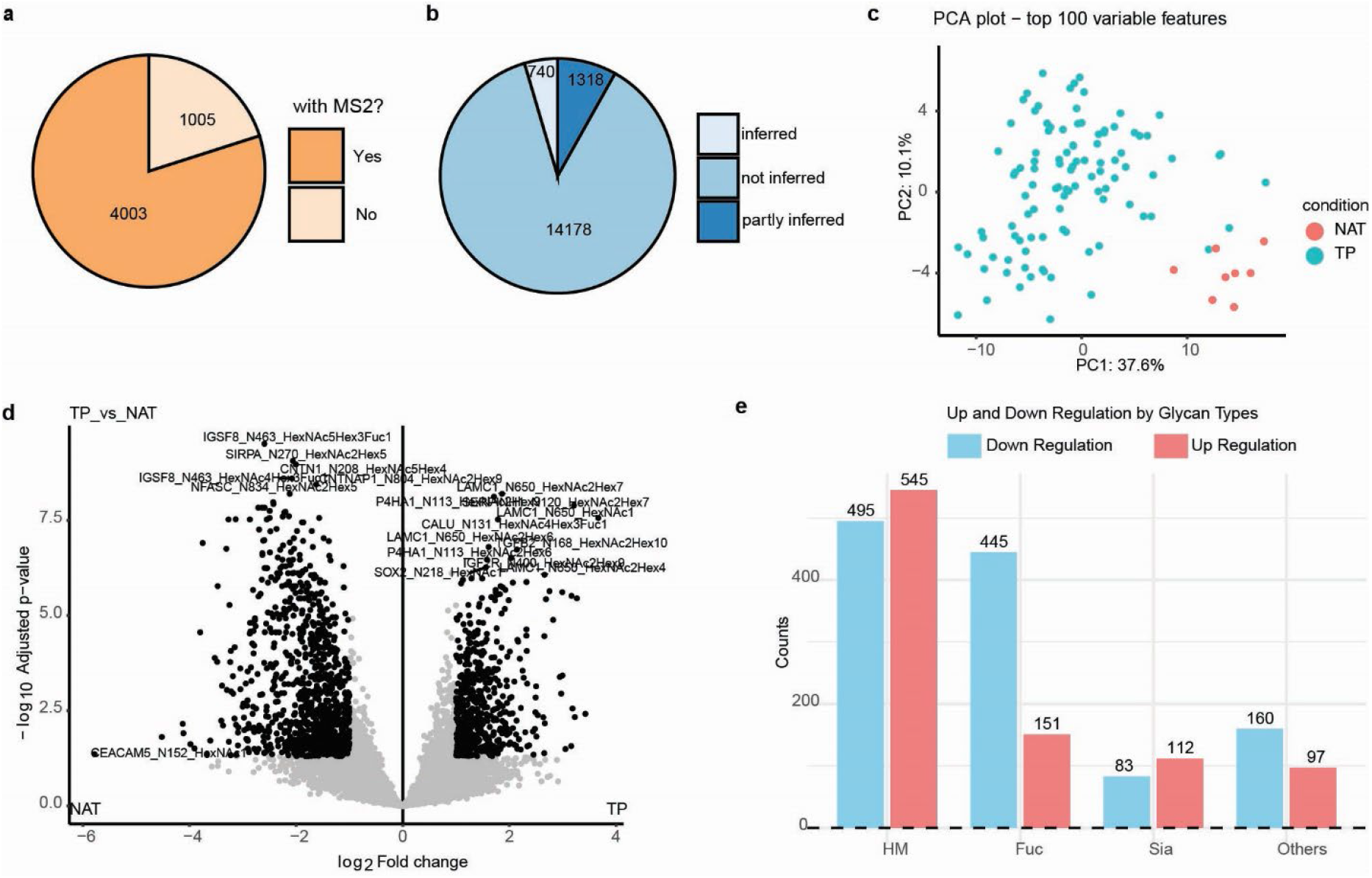
Quantitative glycoproteomic analysis enhanced by MBG and FragPipe-Analyst. a) Numbers of MBG inferred peaks with and without a corresponding MS2 hit. b) Number of unique glycopeptides identified in the GBM dataset identified by MSFragger-Glyco (not inferred), exclusively inferred by MBG (partly inferred) and inferred by MBG but also identified by MSFragger-Glyco in other runs (partly inferred). c) Principal Component Analysis (PCA) plot based on the top 100 glycopeptide features, showing clear separation between grade 4 astrocytomas and normal adjacent tissue. d) Volcano plot depicting differentially expressed glycopeptides (>1.5-fold change). e) Distribution of differentially expressed glycopeptides based on glycan type.

We then performed downstream analysis using FragPipe-Analyst. A PCA plot based on the top 100 features revealed a clear separation between grade 4 astrocytomas and normal brain tissue (**Fig. 4c**). Differential analysis identified 868 upregulated and 1,133 downregulated glycopeptides (>1.5-fold change), including 27 exclusively identified with MBG (Fig. 4d). When categorizing these differentially expressed glycopeptides by glycan type, we observed that fucosylated glycopeptides exhibited a strong tendency toward downregulation, accounting for 70.9% of all fucosylated fold-change glycopeptides (Fig. 4e), consistent with previous observations of altered fucosylation in glioblastoma.

### Expanding identification with adduct or modifications on glycopeptides

Electrospray-ionized glycopeptides often exhibit adduct ions, such as ammonium or metal cations, which have been widely reported in glycoproteomics studies^10,11,17,20^. Additionally, glycan modifications like mannose-6-phosphate (M6P) on high mannose glycans and *O*-acetylation of sialic acids are also observed. However, many of these adducts and modifications typically occur at lower abundance. Including them in database searches can significantly increase the search time and database size, which reduces the sensitivity of searches at a given FDR. MBG effectively identifies these adducts and modifications without requiring their inclusion in the glycan database, allowing them to be recovered without disrupting the search. In cases with more abundant adduction, MBG can also function as a screening tool, where any abundantly detected adducts or modifications in MBG can be added to the primary search.

To evaluate the performance of MBG in identifying adducts, we compared its search time with that of a traditional database search using the pGlyco2 mouse liver dataset, focusing on the identification of three adducts: NH_4_^+^, Fe^3+^, and Na^+^. This dataset was selected because preliminary manual inspection during method development revealed multiple glycopeptides exhibiting spectral features consistent with iron adduction, making it a suitable test case for evaluating adduct identification. A search with adducts added by MBG, but not included in the database search, completed in 21.7 minutes, whereas the traditional method considering the various adducted glycans in the primary search took 115.9 minutes for five runs. Since the presence of adduct ions in this dataset was not previously known, we conducted a straightforward analysis to estimate the prevalence of adduct-containing glycoPSMs by searching for characteristic B ions, specifically HexNAc(1)Hex(1), with Na^+^ or Fe^3+^ in MS/MS spectra. Among 226,702 MS2 spectra examined, 26.0% contained HexNAc(1)Hex(1)+Fe^3+^, while only 0.6% featured HexNAc(1)Hex(1)+Na^+^ (**Table S1**). Given this low frequency, we expect MBG to obtain few Na^+^ matches, and any without the characteristic B ions could be interpreted as entrapment adducts arising from random matches.

The MSFragger-Glyco search without adducts led to 4,386 unique glycopeptides. In addition, MBG identified 1,234 unique glycopeptides, of which 457 were non-adduct glycoforms and 777 carried adducts, including 388 NH_4_^+^, 386 Fe^3+^, and 3 Na^+^ glycopeptides (**Fig. 5a**). By comparing retention time (RT) shifts relative to their parent glycopeptides, the median shifts of NH^4+^ and Fe^3+^ were close to zero, consistent with our expectations for these small cationic adducts (**Fig. 5b**). Since glycan fragment B-ions provide essential compositional information, we further examined inferred glycopeptides alongside their corresponding MS2 spectra. We assessed the presence of characteristic B ions in spectra of Fe^3+^- and Na^+^-adducted glycoPSMs. Excluding those without MS2 hits, MBG identified 685 Fe^3+^-adducted glycoPSMs, of which 554 (80.9%) exhibited the HexNAc(1)Hex(1)+Fe^3+^ B ion. In contrast, none of 4 Na^+^-adducted glycoPSMs detected a feature B ion of HexNAc(1)Hex(1)+Na^+^ (**Fig. 5c**). This stark contrast reinforces that these Na^+^-adducted matches are likely false positives. To further validate Fe^3+^ assignments, we examined both MS2 fragmentation and MS1 isotopic patterns of a representative MBG-inferred glycopeptide. Iron has a pronounced mass defect and multiple naturally abundant stable isotopes, most notably ^54^Fe (5.8%) and ^56^Fe (91.7%), which together produce a characteristic isotope envelope distinct from protonated species. Consistent with the natural isotopic distribution of iron, Fe^3+^-adducted glycopeptides displayed characteristic mass-deficient isotope peaks preceding the monoisotopic precursor, while these pre-monoisotopic peaks were absent in the corresponding protonated glycopeptide (**Fig. S3a, b**). Fragmentation spectra further supported these assignments through the presence of Fe^3+^-containing B and Y ions, including the diagnostic HexNAc(1)Hex(1)+Fe^3+^ fragment (**Fig. S3c**). This exercise clearly demonstrates that many unexplained MS1 and MS2 features in (glyco)proteomics data can occur when considering unusual adducts. In this particular case, we hypothesize that the high occurrence of Fe^3+^ adducts in the glycopeptides from the assessed mouse liver dataset has no biological origin, but is caused by the particular experimental set-up used. This hypothesis is backed by the fact that in the Human Plasma PASEF glycopeptide dataset we generated ourselves, barely any glycopeptides were assigned to bare Fe^3+^ ions.

**Figure 5.**
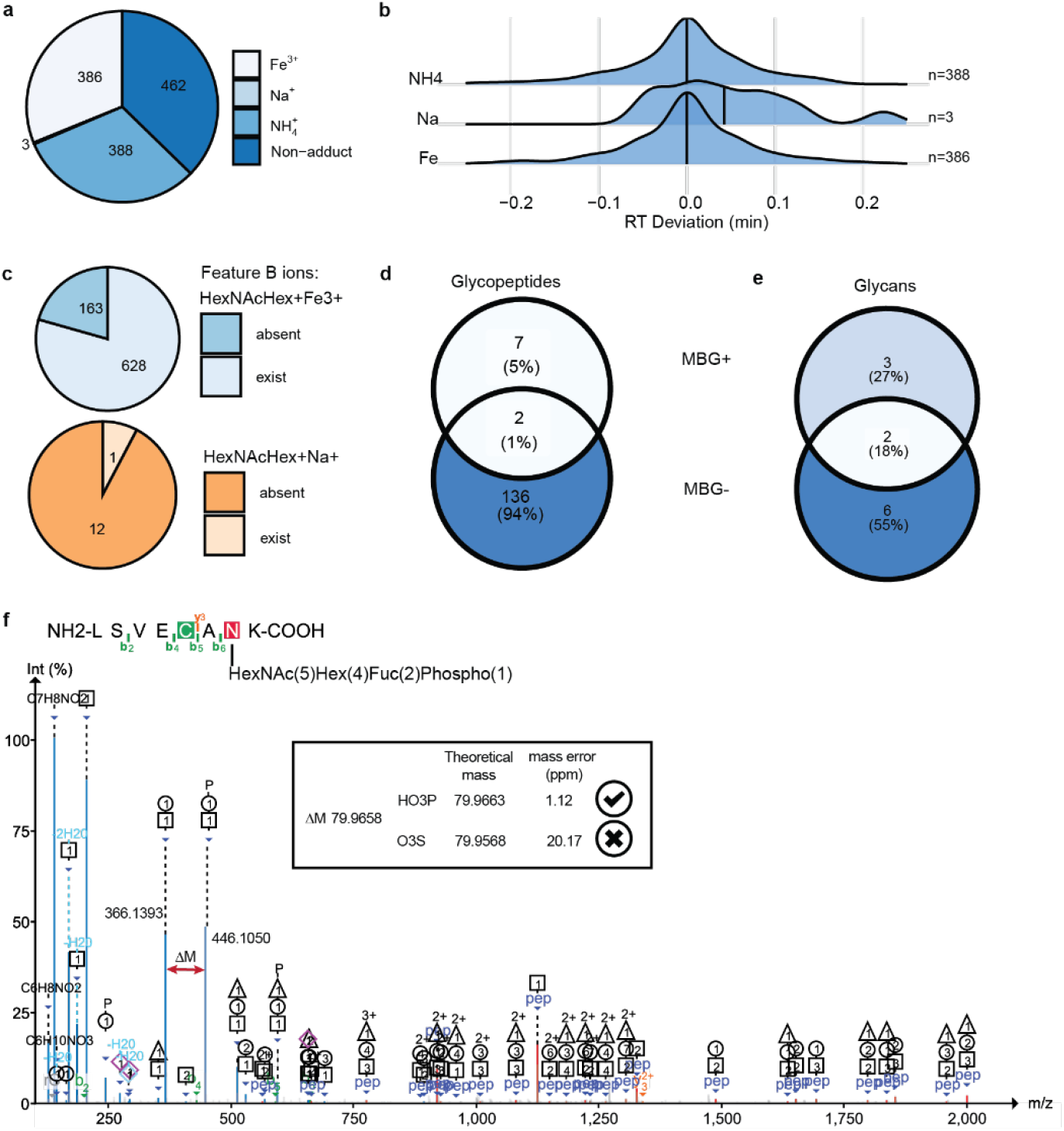
Identification of glycopeptide adducts and glycan modifications using MBG. (a) Pie chart depicting the distribution of 1,234 glycopeptides identified by MBG from mouse liver, including 457 non-adduct glycopeptides and 777 adduct-containing glycopeptides (388 NH4^+^, 386 Fe^3+^, and 3 Na^+^). (b) Retention time (RT) shift comparison of NH4^+^ and Fe^3+^ glycopeptides relative to their parent glycopeptides. (c) Validation of MBG-inferred Fe3+ and Na+ glycoPSMs by the presence of feature B ions. Upper: numbers of Fe3+ glycoPSMs that exhibit or lack the feature B ion HexNAcHex+Fe3+. Bottom: numbers of Na+ glycoPSMs that exhibit or lack the feature B ion HexNAcHex+Na+. (d-e) Identification of phospho-glycopeptides in mouse brain samples: phospho-glycopeptides (d) and glycans (e) identified by MSFragger-Glyco and MBG. (f) MS2 spectrum of glycopeptide LSVECANK with glycan composition HexNAc(5)Hex(4)Fuc(2)Phospho(1), spectrum annotated with FragPipe-PDV.

To further evaluate MBG’s performance in identifying glycan modifications, we focused on detecting mannose-6-phosphate (M6P) glycans in the data obtained from the mouse brain tissue, where they have been well-characterized and relatively abundant. Because phosphorylation typically causes a significant retention time delay, we conducted a search using a customized database containing 14 known M6P glycans. Across five raw files, MSFragger-Glyco identified 89 glycopeptides containing at least one phosphate group, whereas MBG increased this number to 98 (**Fig. 5d**). In addition, MBG inferred 4 phosphate-containing glycan compositions that were not present in the original search database (**Fig. 5e, Table S2**).

To assess the confidence of these identifications, we examined the MS2 spectra of phosphate-containing glycopeptides identified by MBG. Among 17 spectra analyzed, 11 were assigned to canonical M6P glycan compositions, all of which exhibited the characteristic B ion Hex(1)Phosphate(1), a known diagnostic ion for M6P structures. The remaining 6 spectra were assigned to complex type glycans— specifically, HexNAc(4)Hex(4)Fuc(2)Phospho(1) or HexNAc(5)Hex(4)Fuc(2)Phospho(1)—which contain fucose branches. For these cases, we searched for two additional diagnostic B ions: HexNAc(1)Hex(1)Phospho(1) and HexNAc(1)Hex(1)Fuc(1)Phospho(1), which would be expected if the phosphate group resides on a fucosylated branch. Remarkably, 5 out of 6 spectra exhibited both diagnostic ions, supporting the accuracy of MBG’s inference of these complex type phosphorylated glycans. Phosphorylation of complex-type glycans has not, to our knowledge, been previously reported. However, sulfation represents a plausible alternative modification with a similar mass. Therefore, we also performed a stringent mass accuracy analysis of diagnostic fragment ions. **Fig. 5f** shows one representative MS2 spectrum, visualized using PDV-Viewer, the measured mass difference between HexNAc(1)Hex(1)Phospho(1) and HexNAc(1)Hex(1) was 79.9658 Da, corresponding to a mass error of 1.12 ppm relative to the theoretical phosphate mass, compared to a 20.17 ppm error relative to the theoretical sulfate mass, which does support phosphorylation rather than sulfation as the underlying modification.

## Conclusions

We introduce MBG, a modular and efficient add-on method that expands glycopeptide identification using MS1-level inference, without disrupting existing glycoproteomics workflows. By leveraging consistent retention time (RT) and ion mobility (IM) shifts associated with glycan composition changes, MBG enables the inference of “missing” glycoforms that elude conventional searches that consider only a single MS2 spectrum at a time. Since MBG calculates RT and IM shifts separately for each MS file, it can accurately infer glycopeptides within each run regardless of the gradient used in other files. As long as a continuous LC gradient is present within a single MS file, MBG operates effectively, making it fully compatible with fractionated samples or datasets generated with differing gradients across runs. Through integration with the FragPipe suite, MBG facilitates downstream statistical and biological analyses across both label-free and isobaric datasets.

Our benchmarking on a well-characterized fission yeast dataset demonstrated that RT shifts between high-mannose N-glycans are highly consistent across peptides, providing a reliable foundation for inference. Using these shifts, MBG recovered glycopeptides that were not initially identified by MSFragger-Glyco, with a 23.6% increase in unique identifications at a 5% FDR threshold. Importantly, over half of MBG-inferred glycopeptides were supported by corresponding MS2 spectra, and spectral similarity analyses confirmed that inferred glycopeptides shared significantly greater resemblance to their parent spectra than random controls. An entrapment analysis further validated the specificity of MBG, with a false discovery rate of 0.6% for biologically implausible glycans.

We also demonstrated MBG’s scalability and applicability across diverse datasets. In the human plasma PASEF dataset, MBG expanded glycopeptide identification by 14.6%, with most inferred glycans matching known complex fucosylated and sialylated structures. Aberrant fucosylation and sialylation on plasma proteins are known biomarkers in multiple diseases, even in low concentration, making these species clinically important to monitor. Furthermore, by recovering numerous low abundance glycopeptides, MBG contributes to achieving a more complete landscape of glycosylation at each glycosite. Similarly, when applied to a TMT-labeled GBM dataset, MBG identified 740 additional glycopeptides and recovered differentially expressed features that revealed clear biological stratification and glycan-type-specific regulation. These results highlight MBG’s utility for both discovery and comparative glycoproteomics.

Beyond glycoform inference, MBG offers a robust framework for the identification of glycopeptides with adducts or modifications that are typically excluded from database searches due to their low abundance or ambiguous chemical nature. We showed that MBG efficiently identified NH_4_^+^ and Fe^3+^ adducts, while avoiding unexpected sodium adducts, with minimal computational overhead compared to traditional open searching. In a targeted search for phospho-modified glycans, MBG successfully inferred additional glycopeptides with phosphate groups—modifications that are rarely captured in standard analyses but may play significant biological roles.

While these results are promising, several limitations remain. The accuracy of RT and IM inference depends on the density and quality of observed glycan transitions within a dataset. RT shifts can vary over the chromatographic gradient—particularly for certain glycans such as sialic acid-containing species—and can also be influenced by glycan isomerism, which may reduce RT prediction accuracy. Additionally, mobile phase composition (e.g., FA vs. DFA or TFA), stationary phase chemistry, and gradient profiles can all affect glycan elution behavior. Additionally, although MBG includes a decoy strategy for FDR estimation, future work may benefit from incorporating orthogonal validation strategies, such as retention time prediction models trained on glycan-specific features. Finally, MBG currently treats glycan shifts independently and does not account for cases with complex structural isomers—areas that could be addressed by creating and mapping multiple MS1 profiles for single glycopeptide.

In summary, MBG provides a lightweight, data-driven solution for extending glycopeptide coverage using predictable physicochemical behavior of glycan transitions. As glycoproteomics continues to expand toward more comprehensive and systems-level analyses, MBG represents a valuable addition to the computational toolbox, helping to close the gap between observed and expected glycoproteome coverage.

## Conflict of Interest Statement

A.I.N. is the Founder of Fragmatics and serves on the scientific advisory boards of Protai Bio and Infinitopes. A.I.N., D.P., and F.Y. have a financial interest due to the licensing of MSFragger and IonQuant to commercial entities. Other authors have no conflicts of interest.

## Acknowledgements

This work was supported in part by the National Institutes of Health grants R01-GM-094231 and U24-CA271037 (to A.I.N). This research received funding through the Dutch Research Council (NWO) funding the X-omics Road Map program (project 184.034.019).

## Data Availability

All raw data except for the human plasma PASEF data can be found in the public repositories noted in the Methods section. The human plasma PASEF raw data and all processed results (PSM tables) used in the creation of all figures can be found at PRIDE repository (https://www.ebi.ac.uk/pride/archive/) with identifier PXD074575.

