## Supplemental Information for "Expanding Glycopeptide Identification with Match-Between-Glycans in FragPipe"

| **Table of Contents** | S1 |
|---|---|
| **Table S2.** Comparison of Phospho-containing Glycan Identifications in Mouse Brain | S2 |
| **Figure S1.** Distribution of Parent Glycan Types for Inferred Glycan Differences. | S3 |
| **Figure S2.** Relationship between Peptide Intensity and Inferred Glycan Count. | S4 |
| **Figure S3.** Manual Verification of $Fe^{3+}$ Adducted Glycopeptides | S5 |

**Table S1. MBG expansion of M6P glycan identifications in mouse brain.**
The first column lists phospho-glycans included in the customized search database. The second column lists phospho-glycans directly identified by MSFragger-Glyco. The third column shows phospho-glycans inferred by MBG, including compositions not present in the original database. This comparison demonstrates MBG's ability to expand glycopeptide coverage beyond standard database search results.

| Phospho glycans in database | MSFragger-Glyco identified phosoho glycans | MBG-inferred phospho glycans |
|---|---|---|
| HexNAc(2)Hex(6)Phospho(1) | HexNAc(2)Hex(7)Phospho(1) | HexNAc(2)Hex(7)Phospho(1)NH4(1) |
| HexNAc(2)Hex(3)Phospho(1) | HexNAc(2)Hex(7)Phospho(2) | HexNAc(2)Hex(6)Phospho(1)NH4(1) |
| HexNAc(2)Hex(4)Phospho(1) | HexNAc(2)Hex(6)Phospho(1) | HexNAc(2)Hex(5)Phospho(1) |
| HexNAc(2)Hex(5)Phospho(1) | HexNAc(2)Hex(8)Phospho(1) | HexNAc(4)Hex(4)Fuc(2)Phospho(1) |
| HexNAc(2)Hex(6)Phospho(1) | HexNAc(3)Hex(6)Phospho(1) | HexNAc(5)Hex(4)Fuc(2)Phospho(1) |
| HexNAc(2)Hex(7)Phospho(1) | HexNAc(2)Hex(8)Phospho(2) | HexNAc(2)Hex(7)Phospho(1) |
| HexNAc(2)Hex(8)Phospho(1) | HexNAc(3)Hex(7)Phospho(1) | |
| HexNAc(2)Hex(3)Phospho(2) | HexNAc(2)Hex(5)Phospho(1) | |
| HexNAc(2)Hex(4)Phospho(2) | | |
| HexNAc(2)Hex(5)Phospho(2) | | |
| HexNAc(3)Hex(6)Phospho(1) | | |
| HexNAc(3)Hex(7)Phospho(1) | | |
| HexNAc(2)Hex(6)Phospho(2) | | |
| HexNAc(2)Hex(7)Phospho(2) | | |
| HexNAc(2)Hex(8)Phospho(2) | | |

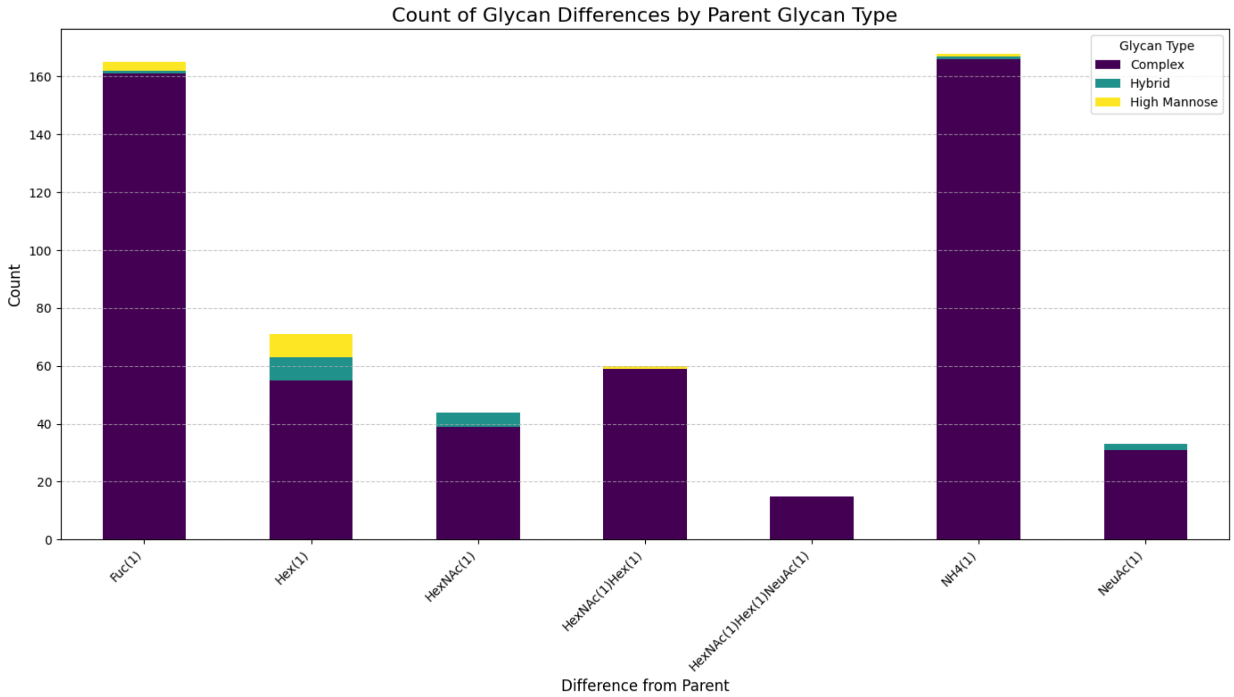


**Figure S1: Distribution of Parent Glycan Types for Inferred Glycan Differences.**

The stacked bar chart shows the total count (y-axis) for each 'Difference from parent' category (x-axis). Each bar is segmented by the classified type of the parent glycan from which the inference was made. Parent glycans were categorized based on their HexNAc count as High Mannose (HexNAc(2)), Hybrid (HexNAc(3)), Complex (HexNAc > 3).

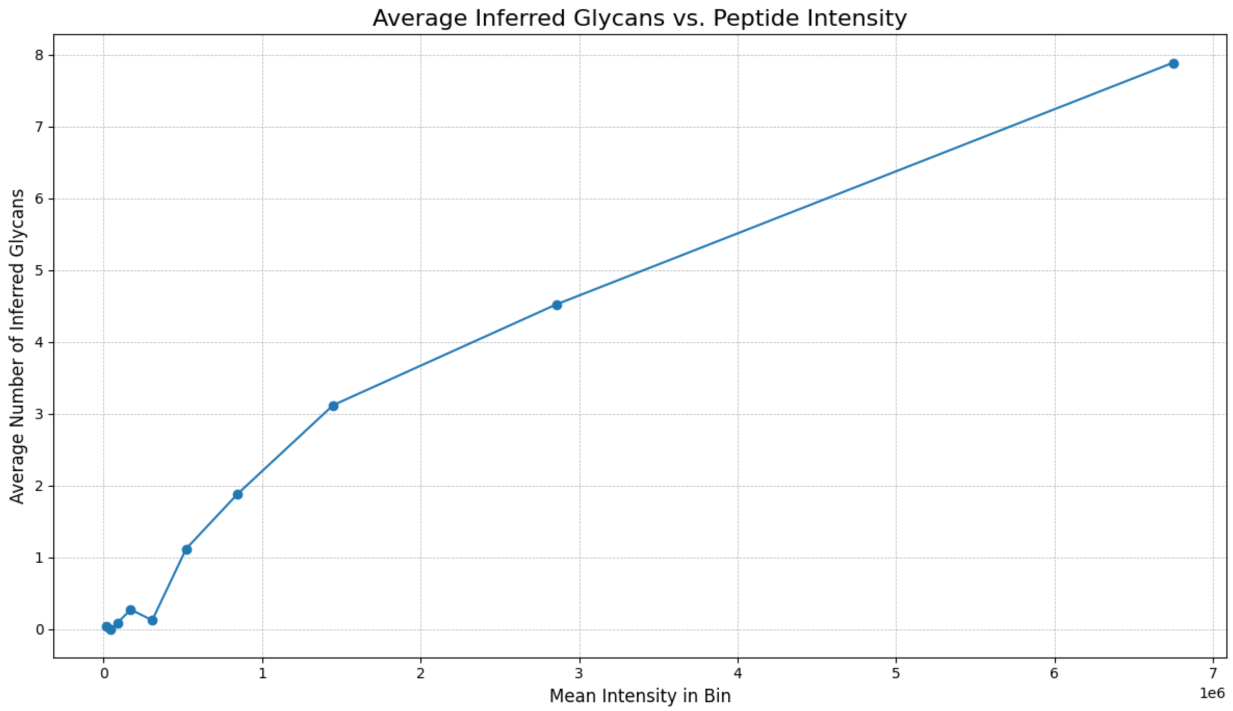


**Figure S2. Relationship between Peptide Intensity and Inferred Glycan Count.**

The plot shows the mean number of inferred glycans (y-axis) as a function of binned peptide intensity (x-axis). The dataset was grouped into 10 quantile bins based on intensity, where each bin contains an equal number of peptides (peptide with intensity=0 not included in this analysis). Each data point represents the average number of MBG inferred glycans and the average intensity for all peptides within that bin. The trend indicates a positive correlation, where peptides with higher intensity are associated with a higher average number of inferred glycans.

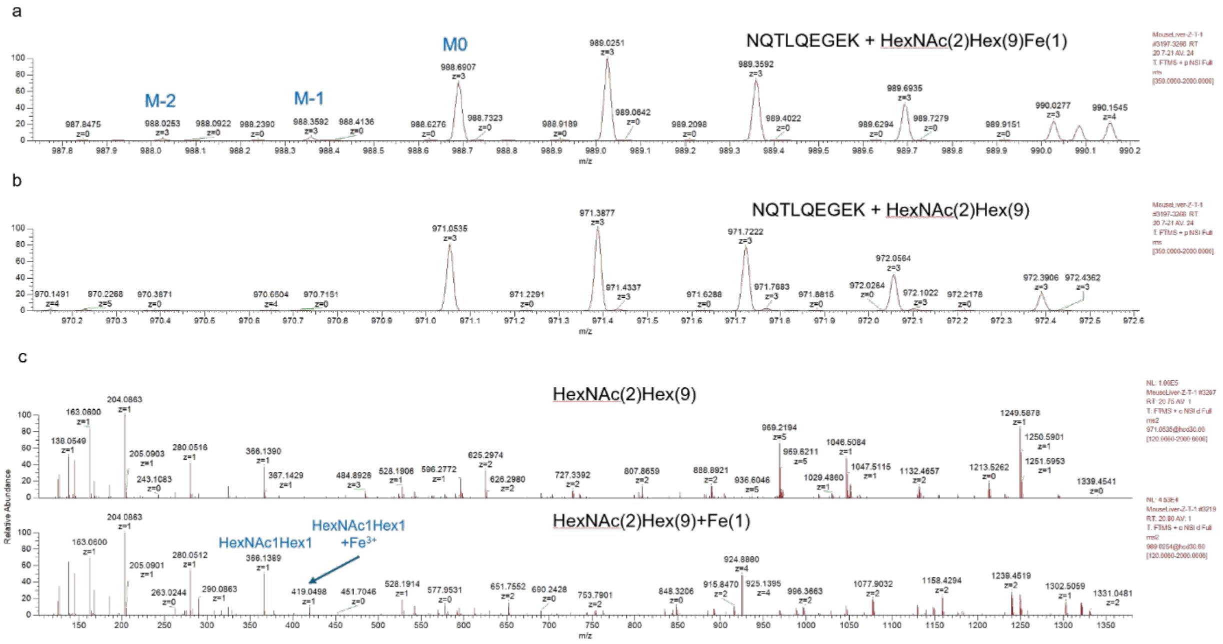


**Figure S3. Manual Verification of $Fe^{3+}$ Adducted Glycopeptides**

Chromatographic profiles of glycopeptide NQTLQEGEK + HexNAc(2)Hex(9) with (a) and without (b) $Fe^{3+}$ adduct. Compared with the profile without $Fe^{3+}$, two low intensity peaks at m/z 988.0253 (M-2) and m/z 988.3592 (M-1) are detected in front of the monoisotopic peak at m/z 988.6907, which is a feature of Fe-containing species. The MS/MS spectra comparison (c) further confirmed the presence of $Fe^{3+}$ adduct, where there are several peaks exhibiting the characteristic +52.91 mass difference (replacement of 3 protons by $Fe^{3+}$) of B and Y ions, including HexNAc(1)Hex(1) + $Fe^{3+}$ at m/z 419.05.
